# Sensory prediction errors predict motor prediction errors

**DOI:** 10.1101/2025.03.16.643175

**Authors:** Dominic M. D. Tran, Nicolas A. McNair, Alexis E. Whitton, Thomas J. Whitford, Evan J. Livesey

## Abstract

When sensory inputs can be predicted by an organism’s own actions or external environmental cues, neural activity is often attenuated compared to sensory inputs that are unpredictable. We have recently demonstrated that attenuation to predictable inputs is also observed when stimulating the motor system with transcranial magnetic stimulation (TMS). Akin to sensory attenuation, predictable TMS excites the motor system less effectively than unpredicted TMS. However, it remains unclear whether these motor prediction signals are related to, or even dependent on, sensory prediction. Using dual-site TMS to target two brain regions, we arranged different warning cues to predict different regions of stimulation and measured motor attenuation using motor-evoked potentials. We found that expecting TMS over the motor cortex produced stronger attenuation than expecting TMS over a non-motor region, confirming that the attenuation observed is directly linked to activation of the motor system and not due to the sensory by-products of TMS. Using combined TMS-EEG, we measured motor attenuation with motor-evoked potentials, and simultaneously measured sensory attenuation to the sound of TMS (a coil “click”) with auditory-evoked potentials. We found that both motor and auditory potentials were attenuated to predictable TMS compared to unpredictable TMS. Critically, the magnitude of auditory attenuation predicted the magnitude of motor attenuation. Our results reveal a close correspondence between error processing in the sensory and motor systems. The findings provide compelling evidence that predictive coding is governed by domain-general properties across distinct neural systems, which share common mechanisms responsible for all forms of predictive learning.

---

Prediction-based attenuation of the brain’s response to sensory inputs is widely observed across different modalities and various organisms. When sensory inputs such as auditory (Mifsud & Whiteford, 2017; Sowman et al., 2012), visual (Tang et al., 2023; Kok et al., 2012), or tactile (Blakemore, Wolpert, & Frith, 1998) information can be predicted by an organism’s own actions or by external warning cues, neural activity is attenuated compared to sensory inputs that are unpredictable. For example, when moving our hand to touch another part of our body, we can predict haptic information from an efference copy of the signal sent to move our hand (e.g., Wolpert et al., 1995). This efference copy allows the tactile sensation to be anticipated in advance of the hand touching the body (i.e., corollary discharge; Sperry, 1950), and results in a less salient sensation being experienced compared to an external source touching the body. This form of sensory attenuation to predictable information is thought to be the reason we are unable to tickle ourselves (Blakemore et al., 1998).

Sensory attenuation is thought to be adaptive from an evolutionary standpoint. Attenuation of predictable or self-generated sensory information allows us to prioritise unexpected and novel sensory information—such as the sound or sight of an incoming predator. Such functions are likely to be fundamental and ubiquitous to the central nervous system given demonstrations of sensory attenuation in many animals, including insects, fish, birds, and mammals (Crapse & Sommer, 2008). Meanwhile dysfunctions in sensory attenuation in humans have also been linked with atypical thoughts and cognitions, with impairments in sensory attenuation, for example, linked with schizotypy traits (Oestreich et al., 2015) and schizophrenia symptoms (Ford et al., 2007).

Modelled in the laboratory, sensory attenuation can be measured using an auditory response task in which a tone is played when the participant presses a key or is played without warning (Ford et al., 2007). Sounds are accompanied by a characteristic auditory-evoked potential when neural activity is measured using EEG. The amplitude of the auditory-evoked N1 component is smaller when sounds are delivered predictably and is larger when sounds are delivered unpredictably (e.g., Harrison, et al., 2021; Mifsud, Beesley, Watson, & Whitford, 2016; Oestreich et al., 2015). Critically, the loudness of these sounds is matched, but it is their predictability that modulates the magnitude of the N1 component. That is, the brain’s neural response to predictable auditory signals is weaker compared to unpredictable auditory signals.

We recently found that prediction-based attenuation is also present in the motor system (Tran et al., 2021; Tran & Livesey, 2021). Using transcranial magnetic stimulation (TMS) we stimulated the left motor cortex and measured motor-evoked potentials (MEP) recorded in the contralateral (right) hand. Across three experiments, we showed that direct stimulation of the motor cortex produced weaker MEPs, or motor attenuation, when the stimulation was predictable compared to when the stimulation was unpredictable (Tran et al., 2021). Notably, in Experiment 2, TMS was delivered under three difference conditions, 1) triggered by the participant (self-generated), 2) triggered shortly after a visual stimulus (warning cue), or 3) triggered with no warning (unpredictable baseline). We showed that MEPs were attenuated in both the predictable conditions (self-generated and warning cue) compared with the unpredictable condition, while the self-generated condition produced stronger attenuation than the warning cue condition. We also found that conditions predicting stimulation 100% of the time resulted in stronger attenuation than conditions predicting stimulation 25% of the time. Notably, the attenuation was observed to a novel event (i.e. TMS) with which most participants had no prior experience; it bypassed sensory channels to directly activate the motor cortex; and did not rely on participants making any motor responses (see Tran et al., 2021, Experiment 3; Tran & Livesey, 2021). This suggests that the brain’s predictive mechanisms are ubiquitous and sufficiently flexible to adapt to completely novel stimuli.

Other neural systems also possess functional characteristics that mirror prediction-based attenuation. For instance, the seminal findings by Shultz and colleagues (1997) can be interpreted as showing that dopamine neurons recorded from the ventral tegmental area of monkeys fired less to predicted delivery of juice compared with unpredicted delivery of juice. Using a cued reward task with rats, Ottenheimer and colleagues (2020) showed that a subset of ventral pallidum neurons encode trial-based prediction errors consistent with the Rescorla and Wagner model of associative learning (1972). These neurons showed greater activity for unexpected delivery of sucrose (following maltodextrin on the previous trial) compared to activity for expected delivery of sucrose (following sucrose on the previous trial). The studies provide evidence that neurons both within and outside of the midbrain show a form of prediction-based attenuation (Shultz et al., 1997; Ottenheimer et al., 2020).

Taking together the evidence of prediction-based attenuation in sensory, motor, and reward systems, it suggests there may be a common mechanism that operates throughout the brain to downweight predictable information and/or upweight unpredictable information. Such a mechanism could be a form of predictive coding that constantly generates mental models of the world from bottom-up information (e.g., Friston, 2005). Unexpected signals that are inconsistent with top-down model predictions—i.e., prediction errors—are important for signalling that the current model does not accurately reflect the environment and requires updating. Attenuating predictable information may be one means for prioritising prediction error signals. Critically, most predictive coding theories are purely grounded in sensory processing, most often visual perception, but our recent results suggest that these predictive coding mechanisms may be domain general and operate using sensory, motor, or reward information in the same way. Few studies have tested this hypothesis, instead studying predictive coding phenomena within one domain.

The current series of studies, we tested the hypothesis that prediction-based neural attenuation is a domain-general property of the brain by comparing prediction errors across domain. First, using dual coil TMS, we provided a novel confirmation that the motor attenuation effect was the direct consequence of predicting motor stimulation and was not caused by predicting the sensory by-products of TMS. Second, using combined TMS-EEG, we simultaneously measured motor and sensory attenuation to assess whether prediction-based attenuation across the motor and sensory systems are linked.

In the first experiment, we used a dual coil setup in which one TMS coil was positioned over the participant’s left motor cortex, and a second control coil was positioned over the right parietal region (approximately over P4 EEG electrode site). The control location was selected because stimulation at this location did not stimulate the motor cortex, did not interfere with visual processing, and could be positioned without touching the motor cortex coil. The aim of this experiment was to determine if expecting stimulation over a control site produced the same degree of attenuation as expecting stimulation over the motor cortex. A key feature of the design is that both coils produce salient sensory by-products during stimulation (an auditory coil “click” and tactile vibration on the scalp), but only one coil reliably stimulated the motor cortex. Hence, if our previous attenuation findings (Tran et al., 2021) are the result of predicting the sensory by-products of TMS, anticipation of the control coil stimulation should also produce MEP attenuation. However, if our previous findings reflect attenuation of the motor system that is generated by predicting stimulation of the motor cortex, anticipation of the motor coil stimulation should produce stronger MEP attenuation. The results test whether MEP attenuation reflects downweighting of neural activity specific to stimulation of motor cortex, or whether MEP attenuation reflects a reaction to the sensory expectation of brain stimulation.

**Figure 1.**
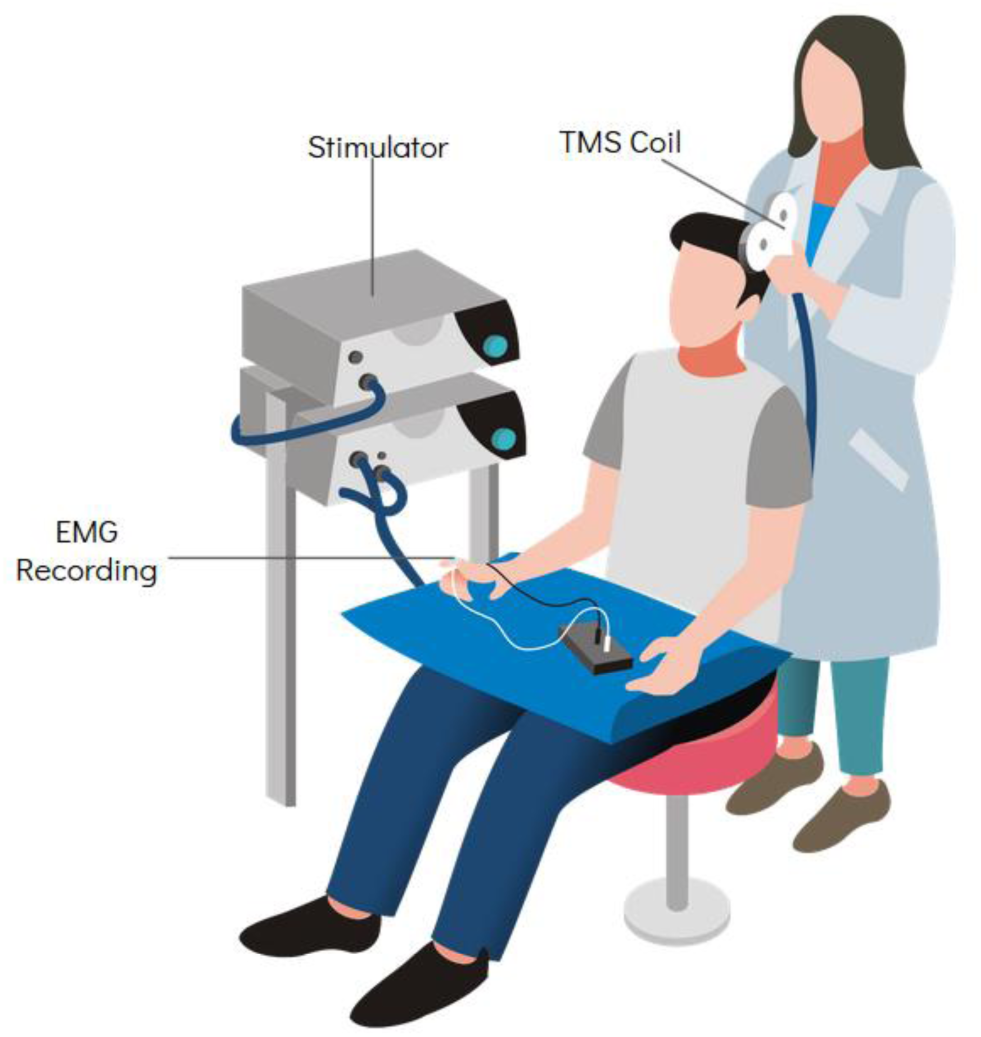
Schematic of the transcranial magnetic stimulation (TMS) setup. Motor-evoked potentials are recorded using electromyography (EMG) attached to the contralateral hand.

In the second experiment, we used a combined TMS-EEG setup to record MEPs from motor cortex stimulation, and auditory-evoked potentials (AEPs) from the coil click when the motor cortex was stimulated. Critically, the TMS stimulation was used to measure attenuation in the motor domain, but the sound generated by the TMS pulse was used as the auditory stimulus to measure attenuation in the sensory domain. The setup therefore allows the simultaneous measurement of motor attenuation to motor cortex stimulation using MEPs and sensory attenuation to the auditory click from the same TMS pulse using AEP. The task was similar to the standard auditory response task used to measure sensory attenuation, but TMS pulses replaced the tones. TMS could be triggered by the participant, followed by a warning cue, or triggered spontaneously without warning. If predictive coding is governed by domain general mechanisms, and these mechanisms vary in strength across individuals, we should see a close correspondence across individuals in the degree to which they show motor and sensory attenuation. That is, we would expect that individuals who show stronger sensory attenuation will also show stronger motor attenuation.

## Results

### Experiment 1

All cues produced attenuated MEPs relative to the baseline condition (see Table 1, left). Replicating our past probability finding (Tran et al., 2021), there was weaker attenuation for the control cue that predicted stimulation of the motor cortex 20% of the time (and nothing 80% of the time) than the cue that predicted stimulation of motor cortex 100% of the time. Critically, the amount of attenuation for the P4 cue was similar to the control cue and weaker than attenuation for the M1 cue. This result suggests that prediction of motor cortex stimulation specifically, rather than prediction of the sensory by-products of TMS, determined the strength of attenuation. Statistically, the M1 cue showed significantly more attenuation than the control cue, *t*(40) = 3.64, *p* = 0.001, *d* = 0.57, and P4 cue, *t*(40) = 2.76, *p* = 0.022, *d* = 0.43. The control cue and the P4 cue were not significantly different, *t*(40) = 0.88, *p* = 1.00, *d* = 0.14, BF_01_ = 3.85. Note, p-values are adjusted for comparing a family of 3 comparisons.

**Figure 2.**
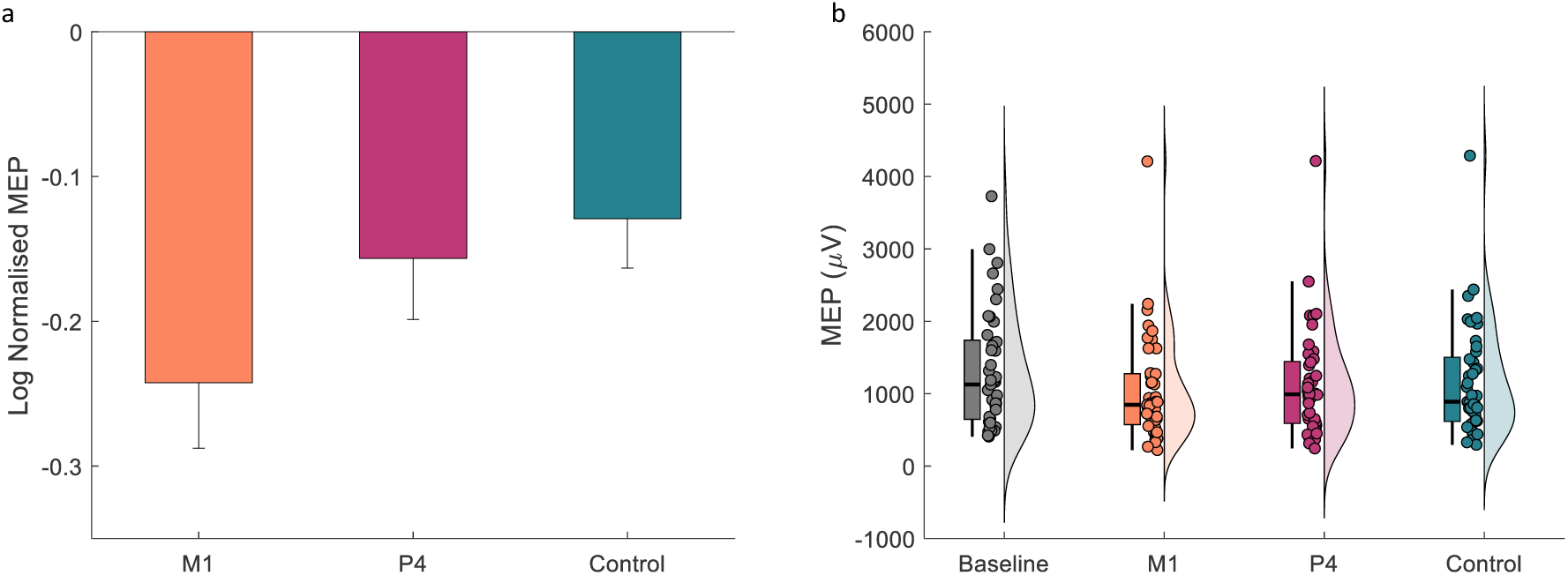
Mean log normalised MEPs to the baseline condition (a) and raw MEPs (b). Error bars represent standard error of the means. MEP = motor-evoked potential; M1 = primary motor cortex location; P4 = P4 10/20 standard EEG electrode location. Note the very positive data points (> 3500 µV) in panel b are from the same participant and do not meet exclusion (> 3SD) when normalising to the Baseline condition for analysis.

**Table 1.** One sample tests of the attenuation effect compared to the baseline condition (H_0_ = 0) for Experiments 1 and 2.

|  | Experiment 1 |  |  | Experiment 2 |  |
| --- | --- | --- | --- | --- | --- |
|  | M1 | P4 | Control | Self | Cued |
| **t** | 5.37 | 3.71 | 3.78 | 4.99 | 6.71 |
| **df** | 40 | 40 | 40 | 34 | 34 |
| **p** | <0.001 | <0.001 | <0.001 | <0.001 | <0.001 |

### Experiment 2

#### MEP

Both the self-generated and warning cue conditions produced attenuated MEPs relative to the baseline condition (see Table 1, right). Replicating our past agency finding (Tran et al., 2021), there was stronger attenuation for the self-generated condition than the warning cue condition, *t*(34) = 2.64, *p* = 0.012, *d* = 0.45, BF_10_ = 3.58.

#### ERP

Based on the peak detection method, the average latency of the N1 peak was, Mean = 104.52 ms, SD = 13.18 ms, and the average latency of the P2 peak was, Mean = 181.73 ms, SD = 24.45 ms. The self-generated and warning cue conditions were attenuated relative to baseline at both the N1 and P2 components. The differences in AUC between the baseline and self-generated conditions were significantly < 0 at N1: *t*(30) = 6.57, *p* < 0.001, *d* = 1.18; and significantly > 0 at P2: *t*(30) = 7.77, *p* < 0.001, *d* = 1.40. The differences in AUC between the baseline and warning cue conditions were numerically smaller in magnitude and significantly < 0 at N1: *t*(30) = 1.90, *p* = 0.033, *d* = 0.34; and significant > 0 at P2: *t*(30) = 2.79, *p* = 0.005, *d* = 0.50.

**Figure 3.**
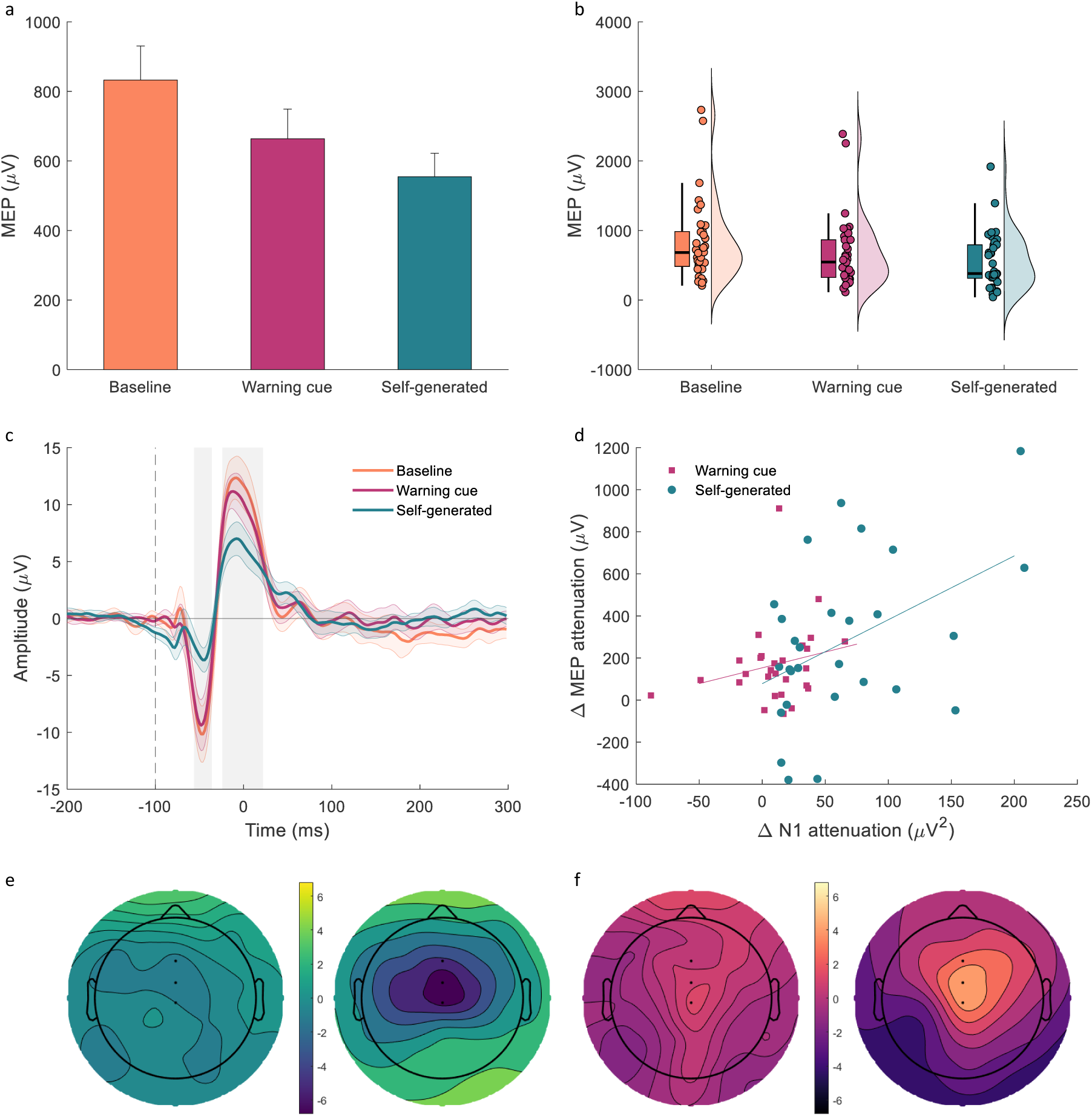
Mean raw MEPs by prediction condition (a) and individual raw MEPs represented as raincloud plots (b). Mean ERPs by prediction condition (c), grey bands represent the N1 and P2 time windows; and scatterplot comparing change in N1 attenuation with change in MEP attenuation (d), delta scores represent different between prediction condition and baseline. Topographic difference maps of the N1 component (e: baseline – warning cue; b: baseline – self-generated) and P2 component (f: baseline – warning cue; baseline – self-generated) with specified electrodes (Fz, FCz, and Cz). MEP = motor-evoked potential; ERP = event-related potential.

#### MEP and ERP relationship

There was a positive relationship between the magnitude of motor attenuation and sensory attenuation at the N1 component. The strength of attenuation of self-generated condition was significantly correlated across MEPs and the N1 component of AEPs, *r* = 0.45, *p* = 0.016, *n* = 28. The strength of attenuation for the warning cue condition was weaker and not significant across MEPs and the N1 component of AEPs, *r* = 0.24, *p* = 0.223, *n* = 28. When aggregating both the self-generated and warning cue conditions, the strength of motor attenuation was positively correlated with the strength of sensory attenuation at N1, *r* = 0.43, *p* < 0.001, *n* = 56.

There was a negative relationship between the magnitude of motor attenuation and sensory attenuation at the P2 component. The strength of attenuation for the self-generated condition was negatively correlated, but not significant, across MEPs and the P2 component of AEPs, *r* = -0.28, *p* = 0.153, *n* = 28. The strength of the attenuation for the warning cue condition was also not significant across MEPs and the P2 component of AEPs, *r* = -0.18, *p* = 0.353, *n* = 28. However, when aggregating both the self-generated and warning cue conditions, the strength of the motor attenuation was negatively correlated with the strength of sensory attenuation at P2, *r* = -0.29, *p* = 0.030, *n* = 56.

## Discussion

The results presented here replicate our initial demonstration that predictable stimulation of the primary motor cortex excites the motor system less effectively compared to unpredictable stimulation (Tran et al., 2021; Tran & Livesey 2021). When participants self-generated TMS or were presented with a warning cue that signalled the imminent arrival of TMS, MEPs in both these predictable conditions were attenuated compared to MEPs triggered by spontaneous, unsignalled TMS. In Experiment 1, we replicated that motor attenuation is sensitive to the strength of the predictive relationship between a warning cue and TMS; the 100% probability cue resulted in greater attenuation than the 20% probability cue. Using dual-coil TMS, we confirmed that the motor attenuation effect is driven by prediction of direct stimulation of the motor cortex, rather than an epiphenomenon caused by predicting the sensory by-products of TMS. In Experiment 2, we found that the typical sensory attenuation effect in AEPs can be replicated using the click sound of a TMS coil that is produced when delivering stimulation. Most noteworthy, we found that when collapsing across predictability conditions (self-generated and warning cue), individual differences in sensory attenuation predicted the strength of motor attenuation, such that individuals who showed greater sensory attenuation at N1 and P2 tended to show greater motor attenuation measured simultaneously in MEPs.

The current findings demonstrate the reliability and robustness of prediction-based motor attenuation measured using TMS. Observing prediction-based attenuation effects in the motor system is particularly interesting because the functional relevance of downweighting predictable motor activity is not immediately obvious. Attenuating predictable sensory activity has clear functional and adaptive benefits as it helps organisms differentiate novel and unexpected information from redundant and repetitive information (Crapse & Sommer, 2008). We have previously hypothesised that prediction-based motor attenuation may function to conserve metabolic energy and enhance signal detection (Tran et al., 2021; Tran & Livesey, 2021). Recent work suggests that task-irrelevant motor movements can overwhelm neural activity (Mathias, 2019; Musall et al., 2019) and these results give credence to the signal detection idea we previously proposed. If organisms can anticipate upcoming motor signals, attenuating these predictable signals can free up the brain to process novel, relevant information. For example, if repetitive motor ticks or habitual actions can be attenuated, any abrupt or spontaneous motor commands can be executed in a more controlled and precise manner. Alternatively, it is possible that attenuating predictable motor signals does not serve a separate adaptive function, but is instead a result of exaptation from existing neural mechanisms controlling functions in other neural systems (e.g., attenuation in the sensory domain). An exciting corollary of this idea is that downweighting predictable activity is a domain-general property of neurons, not specific to the sensory or motor systems.

A number of highly influential frameworks have described sensory processing, particularly visual perception, in terms of predictive coding (Friston, 2005; Rao & Ballard, 1999). The common theme in these frameworks is that the brain is constantly generating controlled, top-down predictions to construct mental models about the external world that are compared with incoming, bottom-up information. Mismatches between these top-down predictions and bottom-up sensory information give rise to prediction errors that signal the mental models require updating. Neural attenuation of predictable information may therefore be a way to increase coding efficiency and maximise sensitivity for detecting prediction errors.

Our current results suggest that predictions based on a mental model and comparisons with incoming information need not be confined to sensory processing, but a similar mechanism can be observed when processing motor information. Supporting this thesis, we found that the magnitude of sensory attenuation at the N1 and P2 components predicted the magnitude of motor attenuation in MEPs. This finding suggests there are variations among healthy individuals in the degree to which prediction modulates neural responses irrespective of the domain in which that information is processed. Moreover, individual differences in neural responsiveness to predictive information has been shown to influence psychological and cognitive function. For example, individuals with schizophrenia do not display the typical N1 suppression effect to predictable sounds (Ford et al., 2007), and schizotypy traits positively correlate with weaker N1 suppression (Oestreich et al., 2015). These impairments in sensory attenuation among individuals with schizophrenia have been hypothesised to account for the presence of delusions of control and even auditory hallucinations, whereby self-generated (predictable) actions or sounds are not sufficiently attenuated and have neural representations that are much more similar to those of externally generated events.

If the capacity to use predictive information to downweight anticipated neural activity varies meaningfully across individuals, and the inability to do so can explain schizophrenia symptomology, deficits in processing predictable information across domains may underlie other clinical disorders. Indeed, impairments in reward prediction error have been reported in individuals with major depressive disorder and bipolar disorder (Whitton et al., 2015). For example, depression has been found to attenuate dopaminergic prediction error signals in a reinforcement learning task (Kumar et al., 2008; Reinen et al., 2021). Building on the idea that predictive coding may be domain-general, the purported deficits in reward prediction error among individuals with mood disorders may be related to deficits in prediction error processing more generally, which should also be observable in the sensory or motor systems. Similarly, dysfunctions in sensory attenuation, for example in patients with schizophrenia (Ford et al., 2007), may reflect a global prediction deficit rather than a domain-specific sensory deficit. This instantiation of predictive coding has important implications for understanding the origins of transdiagnostic and co-occurring symptoms in populations with mental health conditions. Previous researchers have also argued for a general prediction deficit over a domain-specific deficit as a basis for hallucinations and psychosis (Corlett et al., 2019; Powers et al., 2017; Sterzer et al., 2018). However, to our knowledge, no published work has directly tested these ideas by examining how predictions across different systems relate to each other.

Using dual-coil TMS and combined TMS-EEG we were able to isolate the motor attenuation effects to direct stimulation and link attenuation effects across sensory and motor domains. However, both techniques are relatively novel, so it is worth discussing whether the protocols are working as intended, particularly with TMS-EEG. Combining TMS-EEG allowed us to measure AEPs to the sound accompanying TMS stimulation. However, combined TMS-EEG is often used for measuring TMS-evoked potentials (TEPs)—cascades of neural activity resulting from stimulating the brain (Ilmoniemi & Kičić, 2010). For our purposes, it is important to establish, firstly, that the TMS coil click can generate a clean auditory evoked potential with distinct N1 and P2 components; and secondly, that the N1 and P2 components are not solely the product of TEPs. Both these considerations were tested by Rocchi and colleagues (2021) who used noise masking to manipulate the effect of the coil sound on the EEG recording. The authors showed that the sound of the TMS coil can elicit a clean auditory-evoked potential with a negative peak around 100 ms (N1 component) and a positive peak around 200 ms (P2 component). They also found that early activity between 15-65ms was largely due to TEPs, while middle activity between 65-120ms around N1 was largely due to AEPs, and late activity between 120-270ms around P2 was entirely due to AEPs. Therefore, we can be confident that our N1 and P2 components reflect neural activity in response to the sound of the coil click rather than a neural activity from brain stimulation.

In summary, our results show that prediction errors in the motor system can be measured with TMS. The results provide further evidence that prediction-based attenuation can be observed across different neural systems, in sensory, reward, and now motor networks. Most interestingly, we showed that the strength of neural attenuation in one domain predicted attenuation in another, with prediction effects in AEPs related to prediction effects in MEPs. These findings provide compelling evidence that attenuating neural activity to predictable or expected information may be a domain-general property of the brain and provide novel insights and testable hypotheses for how predictive coding mechanisms may be affected in clinical disorders.

## Methods

### Design

Experiment 1 used a dual coil setup to examine the effect of stimulation predictability on MEP attenuation across three cue conditions. One cue reliably predicted motor cortex stimulation with 100% probability (M1 cue); one cue predicted stimulation over the control location with 80% probability (P4 cue) and stimulated the motor cortex on the remaining 20% for measurement trials (note that TMS over a non-motor region cannot produce MEPs so to get a measurement of motor attenuation the P4 cue needs to stimulate M1 some of the time); and one cue predicted no stimulation with 80% probability (Control cue) and stimulated the motor cortex on the remaining 20% as a probability control for the P4 cue. The P4 and the Control cues both stimulate the motor cortex 20% of the time, but the P4 cue is always accompanied by stimulation (the rest of the stimulation is over the parietal lobe). If MEP attenuation is the result of motor cortex stimulation, the P4 and control cues should produce equivalent attenuation since both cues stimulate the motor cortex 20% of the time. However, if MEP attenuation is driven by the by-products of TMS, the M1 and P4 cues should produce equivalent attenuation since both cues trigger a TMS pulse on every trial. There was a total of 244 M1 TMS trials (154 M1 cue, 30 P4, 30 C cue, 30 baseline) and 124 P4 TMS trials.

Experiment 2 used a combined TMS-EEG setup to examine the relationship between motor and sensory attenuation across three predictability conditions. One condition asked participants to make a key press following the presentation of a visual cue, doing so triggered the TMS (self-generated); one condition triggered the TMS shortly after the presentation of a different visual cue (warning cue); and one condition triggered the TMS spontaneously without warning (baseline). There were a total of 324 EEG trials across three cue conditions. Half of the EEG trials were accompanied by TMS stimulation and half of the EEG trials involved the associated cues and actions of each condition but without any stimulation. Taking a difference wave between the TMS and no-TMS EEG trials removes any neural activity resulting from the cue presentation or action generation, leaving only the neural activity resulting from the TMS pulse.

### Participants

All participants completed a TMS safety questionnaire based on Rossi et al. (2009) and provided informed consent before starting the experiment. All protocols were approved by the Human Research Ethics Committee of The University of Sydney.

Experiment 1 was conducted on 46 participants with 1 participant withdrawing due to discomfort from the stimulation. Based on an estimated medium effect size of d = 0.5 (Tran et al., 2021; Experiment 2, probability effect), 34 participants are needed to achieve 80% power for detecting a within participant effect of probability (difference in attenuation between the high predictability M1 cue and the low predictability C cue). We aimed to collect 45 participants to have approximately 40 participants remaining after exclusion since the effect of coil (difference in attenuation between the M1 cue and the P4 cue) could be smaller than the effect of probability. After excluding 4 participants who failed the manipulation check (see Procedure), there were 41 participants remaining for the analysis.

Experiment 2 was conducted on 40 participants. Based on an estimated large effect size of d = 0.8 (Tran et al., 2021; Experiment 2, agency effect), 15 participants are needed to achieve 80% for detecting a within participant effect of prediction cue type. Based on a converted large effect size of ρ = 0.5, 26 participants are needed to achieve 80% for detecting a correlation between attenuation types. One participant had EEG recorded at an incorrect sampling rate and their data were not analysed; 39 participants remained for analysis of either the MEP or ERP data. Four participants were excluded from the MEP analysis because MEPs could not be reliably elicited after increasing the stimulator up to a comfortable intensity for the participant; due to the combined TMS-EEG setup, the TMS coil was further away from the scalp compared to a standard TMS experiment and therefore the average intensity was higher than a typical TMS experiment. Nine participants were excluded from the ERP analysis (see EEG Pre-processing and Analysis section). After exclusions, 35 participants remained for the MEP analysis, 31 participants remained for the ERP analysis, and 28 participants had both MEP and ERP data.

### Apparatus and stimuli

Experiments were run on a Windows 7 PC using PsychoPy3 to control stimulus presentation. Stimuli were presented on a 24-inch monitor (1920 × 1080 resolution, 60 Hz refresh rate) at a viewing distance of ∼ 57 cm. The stimuli presented on screen subtended approximately 2° of visual angle.

#### EMG

Three electrodes were attached to the right-hand for electromyography (EMG) recording. In preparation for recording, the skin was exfoliated with a small scouring pad and wiped with 70% v/v isopropyl alcohol. Two 10-mm diameter Ag/AgCl electrodes were placed in a belly tendon arrangement over the first dorsal interosseus (FDI) muscle to measure MEPs. A ground electrode was placed over the ulnar styloid process of the wrist. EMG activity was recorded from 100 ms pre-stimulation to 400 ms post-stimulation. This signal was digitally converted (sampling rate: 4 kHz, bandpass filter: 0.5 Hz to 2 kHz, mains filter: 50 Hz, and anti-aliasing) and stored offline for analysis using LabChart software (Version 8, ADInstruments).

#### TMS

TMS was administered using a MagStim 200^2^ stimulator and a 70 mm D70^2^ figure-eight coil. Participants wore an elastic cap marked with the 10/20 EEG electrode positions to help locate the hand region of the motor cortex. The coil was held tangentially to the scalp with the coil oriented 45° from the midline. The motor cortex “hotspot” was located by starting from a position 5 cm lateral and 1 cm anterior to Cz. The coil was then moved around until the maximal MEP could be elicited in the FDI. Once the hotspot was determined and marked, the participant was asked to place their head on a chin and forehead rest for the coil to be locked in position with an adjustable mechanical arm (Manfrotto). Resting motor threshold (rMT) was defined as the lowest stimulation intensity that produced an MEP of at least 50 µV in 5 out of 10 consecutive trials (Rossini et al., 2015).

During the experiments, the stimulation intensity was set to 120% of rMT unless the participant requested a lower intensity due to discomfort (usually 110% of rMT). Participants were asked to keep their head still during thresholding and while the experiment was in-session, but they could move any other time. Since both experiments used a within-participant design where all conditions were experienced by each participant, there was no need for sham TMS. Any peripheral effects of the TMS (tactile and auditory) are matched across all conditions.

For the dual coil set-up (Experiment 1), a second 70 mm figure-eight coil was positioned approximately over the location of the P4 electrode under the 10/20 positioning standards. This coil was connected to a second MagStim 200^2^ stimulator and the stimulation intensity was set to the same value as the first stimulator. The mean stimulation intensity from all participants who started the experiment was M = 52.78% of the maximum stimulator output, SD = 11.37 (n = 46).

For the combined TMS-EEG set-up (Experiment 2), the TMS coil was covered with a 10 mm layer of pliable foam to minimise vibration of the coil on the EEG electrodes. The mean stimulation intensity from all participants who started the experiment was M = 75.15, SD = 12.29 (n = 40). Note that the mean stimulation intensity is higher compared to Experiment 1 due to the extra distance separating the coil and the scalp due to the addition of the foam and EEG electrodes.

#### EEG

In Experiment 2, EEG data were sampled continuously at 10,000 Hz from 64 Ag/AgCl active electrodes (actiCHamp, Brain Products, Gmbh, Gilching, Germany). The electrodes were situated in an EasyCap (GmbH, Woerthsee-Etterschlag, Germany) that positioned them approximately in standard 10/20 locations. Impedances were kept below 10kΩ. Data were acquired using electrode FCz as the online reference and then later re-referenced offline to the average of all 64 scalp electrodes.

### Procedure

#### Experiment 1

Participants passively viewed shapes presented on the computer screen and did not need to make any responses. Three shapes (+, o, and = characters) were randomly allocated to serve as one of the three different prediction cues. The M1 cue was followed by stimulation of the M1 coil 100% of the time, the P4 cue was followed by stimulation of the P4 coil 80% of the time and M1 coil 20% of the time, and the control cue was followed by nothing 80% of the time and stimulation of the M1 coil 20% of the time. The assignment of shapes to cues were counterbalanced between participants. Each cue was presented for 1200 ms, and TMS was triggered after 800 ms. Presentations of the cues were pseudorandomised such that no single cue could appear more than five times in a row. Cues were separated by a variable intertrial interval (ITI) between 2000-3000 ms. Baseline TMS trials (unpredictable condition) occurred between two ITI periods with no change to the screen presentation. At the beginning of the experiment participants were instructed to pay careful attention to the shapes that appeared because they would be asked some questions at the end of the experiment. The questions acted as our attentional manipulation check since there were no response requirements to check for non-compliance. If participants were unable to identify the relationship between the shapes and the location of stimulation, they were excluded from the data analysis. Piloting suggested that this relationship should be easily detected if participants were attentive during the task. Four participants could not identify the relationship of any of the three shapes, and their data were not included in the analysis.

#### Experiment 2

Two shapes (+ and o presented as in Experiment 1) were randomly allocated to serve as cues to signal the two different prediction conditions: the self-generated cue remained on the screen until participants pressed the space key with their left thumb, and TMS was triggered immediately for 50% of these key presses (such that there were equal numbers of self-generated TMS and self-generated no-TMS trials); the warning cue remained on the screen for 1500 ms, and TMS was triggered after 50% of the warning cues (such that there were equal numbers of warning cue TMS and warning cue no-TMS trials). On trials where TMS followed the warning cue, stimulation was triggered after a duration equal to the median reaction time sampled from the self-generated condition; if no reaction time had yet been sampled, the delay was set to 500 ms until a response was recorded. Presentation of the cues were pseudorandomised such that no single cue could appear more than four times in a row. Cues were separated by a variable ITI between 2000-3000 ms. Baseline trials occurred between two ITI periods with no change to the screen presentation; TMS was triggered on 50% of the baseline trials (unpredictable condition) and the remaining 50% of trials were used as no-TMS control trials.

### Pre-processing and Analysis

#### MEP

MEP pre-processing was performed using custom software written in Python (github.com/nicolasmcnair/MEPAnalysis). Trials were excluded if EMG activity pre-stimulation had either an amplitude that exceeded 50 µV, or a root mean square power greater than 3 standard deviations above the mean of all trials for that participant. Mean MEP amplitudes by condition were calculated from the remaining trials using peak-to-peak difference values. Mean MEP data for each condition were log-normalised to the baseline condition at a participant level (Tran et al., 2021). Log-normalised values significantly different from zero indicate an attenuation effect compared to baseline. Log-normalised values significantly different from each other indicate a condition effect.

#### EEG

For Experiment 2, EEG pre-processing was performed using EEGLAB toolbox (v2022.0; Delorme & Makeig, 2004) in MATLAB (R2022a, MathWorks, Natick, MA). Data encompassing TMS pulse artefacts were first replaced using temporal interpolation via ARfit Studio (Schneider & Neumaier, 2001). A learning window from -100 to -5 ms of the TMS pulse was used to inform an autoregressive model. This was then used to replace the data in a window starting at -5 ms up until 45 ms (determined on an individual participant basis). Finally, the original data was blended with the model over an additional period of 5 to 10 ms for all participants. The overall interpolation window was selected on a participant basis to be as short as possible. The primary event of interest was the N1 component of the AEP, where the strength of sensory attenuation has been linked to schizophrenia (Ford et al., 2007; Oestreich et al., 2015). Studies have also found sensory attenuation reflected in the P2 component in healthy adults, but its magnitude is not predictive of schizophrenia. Therefore, we analysed P2 as a secondary event of interest. Both these components were unaffected by the interpolation window that removed the TMS pulse artefact. The purpose of removing the artefact was for visualisation purposes to scale the N1 and P2 components within an appropriate range. The data were then bandpass filtered (0.1 – 30 Hz cutoff frequencies) and downsampled to 250 Hz. Line noise (50 Hz) was removed using the Cleanline function from the PREP Pipeline plugin (Bigdely-Shamlo, Mullen, Kothe, Su, & Robbins, 2015). Bad channels were identified using the Clean Rawdata plugin based on the presence of flatline data (> 5 s), remaining line noise (> 5 SDs compared to signal), or lack of correlation with nearby channels (r < .8; Mullen et al., 2015). These were then replaced with interpolations derived from the other electrodes, weighted by distance. Noisy sections of data were cleaned using artifact subspace reconstruction (ASR) from the Clean Rawdata plugin (Mullen et al., 2015). Data were then re-referenced to the average, before epoching 1000ms prior to and after each event marker—baseline-corrected from -1000 to 0ms of the interpolated waveforms.

Independent components analysis (ICA) was used to identify and remove artefactual components (e.g., eye blinks, muscle activity, etc.) with the assistance of the ICLabel plugin (Pion-Toanchini, Kreutz-Delgado, & Makeig, 2019). The resulting data were imported into ERPLAB (v9.0; Lopez-Calderon & Luck, 2004), and re-epoched from 200ms before to 800ms after each event marker. Finally, individual epochs were removed if they exhibited a peak-to-peak amplitude difference of larger than 100μV in a 200ms moving window, or a sample-to-sample difference of greater than 30μV. Six participants were excluded for having less than half the total number of trials remaining after artefact rejection. Based on a visual inspection of the waveforms and scalp topographies, two participants were found not to generate a clear N1 component and were excluded from the analysis.

Difference waves were created by subtracting the average of the non-TMS trials from the TMS trials for each of the self-generated, warning cue, and baseline conditions. The purpose of subtracting non-TMS trials is to remove any EEG activity that is unrelated to the effect of TMS, this step is most critical for the self-generated and warning cue conditions which involve a key press and/or the onset of a visual stimulus. The N1 component was identified for each individual by detecting the largest negative deflection in a window from 50 – 150 ms post-stimulus onset over electrodes Fz, FCz, and Cz in their grand average ERP waveform (i.e., averaged across all conditions). The boundaries of the component were then identified by finding the two timepoints— earlier and later than the peak—where the amplitude reached half its maximum. For each condition, the area-under the curve (AUC) was then calculated between these two time points. The P2 component was identified by detecting the largest positive deflection in a similar manner between 150 – 250 ms post stimulus.

#### MEP × ERP

To examine the relationship between motor attenuation in MEPs and sensory attenuation in ERPs, delta scores were computed from raw MEPs and AUC of the N1 and P2 components. For the self-generation condition: ΔMEP scores were calculated for each participant as the difference between the mean baseline MEP and the mean self-generation MEP; ΔN1 and ΔP2 scores were calculated for each participant as the difference between the baseline AUC and the self-generation AUC. For the warning cue condition: ΔMEP scores were the difference between the baseline and warning cue MEP values; ΔN1 and ΔP2 scores were the difference between the baseline and warning cue AUC values.

## Author Contributions

D. T., T. W., and E. L. designed research; D. T. performed research; D. T., and N. M. analyzed data. D. T., N. W., A. W., T. W., and E. L. wrote the paper.

## Acknowledgments

The authors thank Julia Lui for their helped with data collection. The research was supported by the Australian Research Council’s Discovery Projects funding scheme (Project Number DP190100410 and DE220100829).

## Notes

### Competing Interest Statement

The authors have declared no competing interest.

